# Population-level deep sequencing reveals the interplay of clonal and sexual reproduction in the fungal wheat pathogen *Zymoseptoria tritici*

**DOI:** 10.1101/2020.07.07.191510

**Authors:** Nikhil Kumar Singh, Emilie Chanclud, Daniel Croll

## Abstract

Pathogens can rapidly surmount crop resistance challenging global food security. On annual crops, pathogens must re-infect from environmental sources at the beginning of every growing season. Fungal pathogens evolved mixed reproductive strategies to cope with the distinct life cycle challenges of colonizing plants, spreading within fields and ultimately producing propagules for survival in winter. However, how genotypic diversity evolves over this period remains largely unknown. Here, we performed a deep hierarchical sampling in a single experimental wheat field infected by the major fungal pathogen *Zymoseptoria tritici*. We analyzed whole genome sequences of 177 isolates collected from twelve distinct cultivars replicated in space at three time points of the growing season. The field population was highly diverse with 37 SNPs per kilobase and a linkage disequilibrium decay within 200-700 bp. We found that ~20% of the individual isolates were grouping into 15 clonal groups. Pairs of clones were disproportionally found at short distance (<5m) but a low degree of dispersal occurred also at the scale of the entire field consistent with a predominant leaf-to-leaf dispersal. We found no association of wheat cultivars and clonal genotypes with the exception of one cultivar. Surprisingly, levels of clonality did not increase over time in the field although reproduction is thought to be exclusively asexual during the growing season. Our study shows that the pathogen establishes vast and stable gene pools in single fields over the growing season. Monitoring short-term evolutionary changes in crop pathogens will inform more durable strategies to contain diseases.

**Data summary:** All Illumina sequencing datasets are available from the NCBI Sequence Read Archive (https://www.ncbi.nlm.nih.gov/sra). The Supplementary Tables S1-S3 list the exact strain names, collection location, genotype and genetic diversity indices.

## Introduction

Infectious diseases have a major influence on the demography of plant and animal populations including humans. Designing effective strategies for disease prevention requires an understanding of the evolutionary dynamics of pathogenicity. A critical factor is the genetic diversity of the pathogen, which strongly impacts disease severity and the spread in heterogeneous host populations (Read & Taylor, 2001; Thrall & Burdon, 2003; Vignuzzi et al., 2006). Host genotypes can vary in their composition of resistance genes, hence pathogens face heterogeneous selection pressures to overcome host resistance. Adaptation to overcome different resistance genes often requires independent mutations (Smith, 1904; Thrall & Burdon, 2003; Alizon *et al.*, 2009). Previous studies have shown that highly resistant host populations harbour the most virulent pathogens, whereas avirulent pathogens dominate susceptible populations (Thrall & Burdon, 2003). Therefore, the evolution of pathogenicity is directly influenced by host diversity. In addition to selection by host genotypes, pathogen populations are subject to selection at life cycle stages outside of the host (Walther & Ewald, 2004; Linde, 2010; Hamelin *et al.*, 2011). Pathogens of annual plants must survive periods of time in the absence of a host and succeed in re-infecting new plants (Laine & Barrès, 2013; Mariette *et al.*, 2016). During the growing season of the host, pathogens are selected to colonize additional plants. Each life cycle stage likely impacts levels of genetic diversity in pathogen populations. However, it remains largely unknown what factors influence spatial and temporal variation in pathogen genetic diversity (Barrett et al., 2008).

In agricultural host-pathogen systems, infections begin with the exposure of a susceptible host to pathogen strains at the beginning of the growing season. The intensity of an infection depends on the level of host resistance, the amount of pathogen inoculum and environmental factors (Burdon *et al.*, 2016; Suffert *et al.*, 2019). If the infection is successful, pathogens complete their life cycle within the host and then transmit to another host. The genetic architecture of traits to successfully passage between infection stages is likely distinct (Jones & Dangl, 2006; Dodds & Thrall, 2009). Entering a host plant requires the manipulation of the host immune system (De Torres *et al.*, 2006; Sun *et al.*, 2006; Nomura *et al.*, 2006). Proliferation and reproduction inside the plant depends likely on optimal resource exploitation and continued suppression of immunity (Chisholm *et al.*, 2006; Steinberg, 2015). Dispersal between hosts is governed by propagule properties (DOBBS, 1942; Peay & Bruns, 2014). As a consequence, pathogen populations likely experience complex selection pressures over the life cycle. Host populations with qualitative resistance will select for pathotypes with matching virulence genotypes (Zhan *et al.*, 2012; Burdon *et al.*, 2016). Research has focused mostly on studying large-effect loci, which encode predominantly functions in pathogen detection (Corwin & Kliebenstein, 2017). In contrast, quantitative host resistance can favour the accumulation of virulence alleles of small, additive effects over longer timescales (Andrivon *et al.*, 2013; Burdon *et al.*, 2016). Therefore, quantitative plant-pathogen interactions can select for continuous distributions of virulence phenotypes in the pathogen population (Corwin & Kliebenstein, 2017). Knowledge of pathogen traits promoting infection at various stages helps to deploy crop varieties with more durable resistance mechanisms. Alternatively, crop rotation and mixed farming can create heterogeneous selection pressure slowing adaptation (Burdon & Thrall, 2008; Stukenbrock & Bataillon, 2012). In any such scenario, pathogens must re-infect crops annually or stably infect perennial host plants, therefore considering the time-scale of pathogen population turnover is important.

Pathogens often have mixed reproductive strategies meaning that sexual and asexual reproduction can alternate over the course of the life cycle adding to the complexity of recognizing species and challenging genetic diversity analyses (McDonald, 1997; Billiard *et al.*, 2012). Fungal crop pathogens often reproduce asexually during the growing season to maximize dispersal at low population density. On senescent plants, sexual reproduction can lead to favourable allele combinations for long-distance dispersal and survival (McDonald & Linde, 2002). An important pathogen with a mixed reproductive strategy is the fungal wheat pathogen *Zymoseptoria tritici*. The fungus is a haploid ascomycete and one of the most destructive pathogens of wheat with yield losses of 5–60% (Fones & Gurr, 2015). Populations worldwide harbour significant variation in pathogenicity and diversity at loci underlying host specificity (Zhong *et al.*, 2017; Hartmann *et al.*, 2017; Hartmann & Croll, 2017; Krishnan *et al.*, 2018). The pathogen follows typically a sequential regime of sexual and asexual reproduction cycles over the growing season. Ascospores are thought to be the source of primary infection in the field and can disperse over long distances (Chen & McDonald, 1996). Once infections are established in a field, *Z. tritici* produces asexual pycnidiospores, which can disperse with rain splash and initiate secondary infections nearby. The spread within fields during the growing season is thought to be predominantly clonal (Chen & McDonald, 1996). Population genomics analyses of worldwide pathogen samples showed high genetic diversity within fields and extensive gene flow (Hartmann *et al.*, 2017, 2018; McDonald *et al.*, 2019). Multi-year studies of *Z. tritici* populations revealed trade-offs between the intra- and interannual scales in the evolution of aggressiveness (Suffert *et al.*, 2018) with re-infections at the beginning of the growing season imposing no detectable genetic bottleneck (Morais *et al.*, 2019). Consistent with the maintenance of high levels of diversity, individual wheat leaves are frequently infected by multiple different genotypes (Linde *et al.*, 2002). However, whole genome analyses of hierarchical field collections to resolve the spatial and temporal changes in genotypic diversity have been lacking.

In this study, we have whole-genome sequenced 177 distinct isolates of *Z. tritici* from a single wheat field covering three time points over the growing season. We analyzed twelve distinct cultivars replicated in space in a randomized block design. Based on genome-wide polymorphisms, we reconstructed how different chromosomal compartments contributed to overall diversity. Using information about the spatial organization in the field and the cultivar of origin, we determined the extent of clonal spread in space and time.

## Results

### Pathogen field population shows extensive sequence variation

A total of 177 isolates of *Z. tritici* were collected from an experimental wheat field in Switzerland planted with 335 wheat cultivars in 1.2-by-1.7m plots (Supplementary Table S1; Karisto *et al.*, 2018). We collected isolates at three time points during the wheat growing season representing different phases in the infection life cycle of the pathogen (Fig 1A; Karisto *et al.*, 2018). The isolates from the first collection (C1) were exposed to the application of demethylation inhibitor (DMI) fungicides. Isolates from the second and third collection were exposed additionally to a succinate dehydrogenase inhibitor (SDHI) and additional DMIs (Fig 1A). We collected isolates from 12 winter wheat cultivars (*n* = 1-43 isolates per cultivar) which were planted in two replicate plots separated by ~100 m (Fig 1B-C). The cultivar Claro was sampled on two additional plots. The collection was performed when the wheat plants were in growth stages GS 41 (C1), GS 75 (C2) and GS 85 (C3), respectively (Meier, 2001; Karisto *et al.*, 2018). We Illumina sequenced whole genomes for all 177 isolates producing 0.8-15 million reads per isolate with an average read count of 4.2 million (Fig 1D). The isolates were sequenced to a depth of 5-77 X and a mean coverage of 21.4 X. We detected a total of 1’496’037 high-confidence SNPs. The average genotyping rate was 97.85 % across loci (Fig 1E). We detected on average 37 SNPs per kb showing that the field population is highly polymorphic. In line with this, we found that the minor allele frequency spectrum showed a strong skew towards rare alleles in the population, which suggests that the population did not experience any recent bottlenecks (Fig 1F). Based on the finding that our population shows substantial genetic variation, we aimed to estimate the total genetic diversity in the field. For this, we performed random down-sampling analysis and we found that subsets of 10% and 50% of the population harboured on average 763’765 and 1’234’846 segregating SNPs representing 51% and 82.5% of the total detected SNPs, respectively (Fig. 1G). We observed only a weak plateau effect for subsets close to 100% indicating that the total polymorphism in the field population is likely substantially higher.

**Figure 1:**
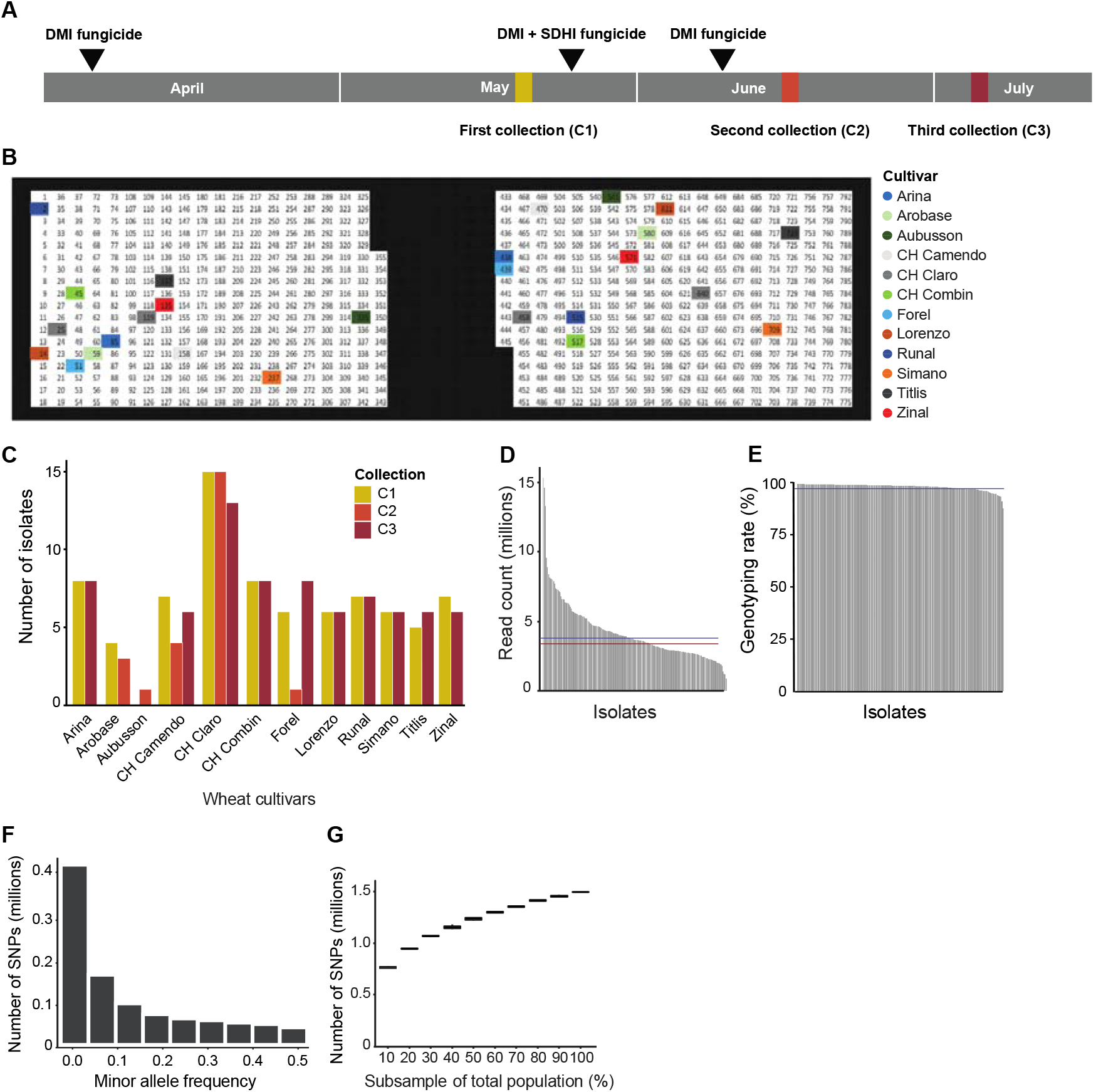
Genome sequencing of 177 *Zymoseptoria tritici* isolates from a single field. A) Time scale showing the three collection points from May to July. The experimental field was treated with fungicides thrice (see Methods for details). B) Graphical representation of the experimental wheat field showing the sampled cultivars (colored background). Each number indicates a plot of 1.2-by-1.7 meters. Wheat cultivar color codes are maintained throughout the manuscript. C) Number of isolates collected from each of the 12 cultivars at each collection time point. D) Trimmed read count for each isolate. Lines represent the mean (blue) and median (red) read counts. E) Genotyping rate of all the 177 isolates (% loci genotyped). The blue line represents the mean genotyping rate. F) Minor allele frequency spectrum of 1’496’037 SNPs genotyped in 177 isolates. G) Boxplot showing the number of segregating SNPs in subsets of the 177 isolates. Error bars show 10 replicates of resampled isolates.

### Accessory chromosome polymorphism

*Z. tritici* harbours the most extensive set of accessory chromosomes known for plant pathogens (Croll & McDonald, 2012). We analyzed chromosome-wide read coverage to identify chromosomal presence-absence variation. All core chromosomes (1-13) were consistently detected in all isolates as expected. In contrast, we found 139 isolates (~79%) missing one or more accessory chromosomes (Fig 2A). We found no isolate that lacked all accessory chromosomes, but we identified two isolates with only a single accessory chromosome out of eight. Accessory chromosome 16 was the most frequent (174/177 isolates) and chromosome 18 was the rarest being present in less than half of the isolates (Fig 2B). We identified several accessory chromosomes showing evidence for partial deletions or duplications based on normalized read coverage (Supplementary figure S1). Specifically, we identified two partial deletions of chromosome 17 and 20 (Fig 2C), as well as chromosomal duplications of chromosome 17, 18 19 and 21 (Fig 2C). In a next step, we analyzed how polymorphism was structured among core and accessory chromosomes. We found that the accessory chromosomes show higher SNP densities but lower total SNP counts compared to core chromosomes (Fig 2D-E). Overall, smaller chromosomes tend to have higher SNP densities (Fig 2F). This is consistent with relaxed selection leading to accumulation of mutations on these chromosomes (Stukenbrock *et al.*, 2010). Another important component of genetic variation is the extent of linkage disequilibrium. We estimated linkage disequilibrium decays for each chromosome separately and found that most accessory chromosomes showed a faster decay compared to core chromosomes (*r^2^* = 0.2 within 100 bp, Fig 2G), with the exception of chromosome 16 and 18 showing a decay of *r^2^* to 0.2 within 300 and 900 bp, respectively, comparable to the decay on core chromosomes (Fig 2H). Interestingly, chromosome 7 showed the slowest decay of all core chromosomes (*r^2^* = 0.2 at ~700 bp, Fig 2H). This chromosome was proposed to have originated at least partially from an accessory chromosome (Schotanus *et al.*, 2015). Taken together, we show that the accessory chromosomes accumulated more sequence variation compared to core chromosome and but that the level of conservation varies substantially among accessory chromosomes.

**Figure 2:**
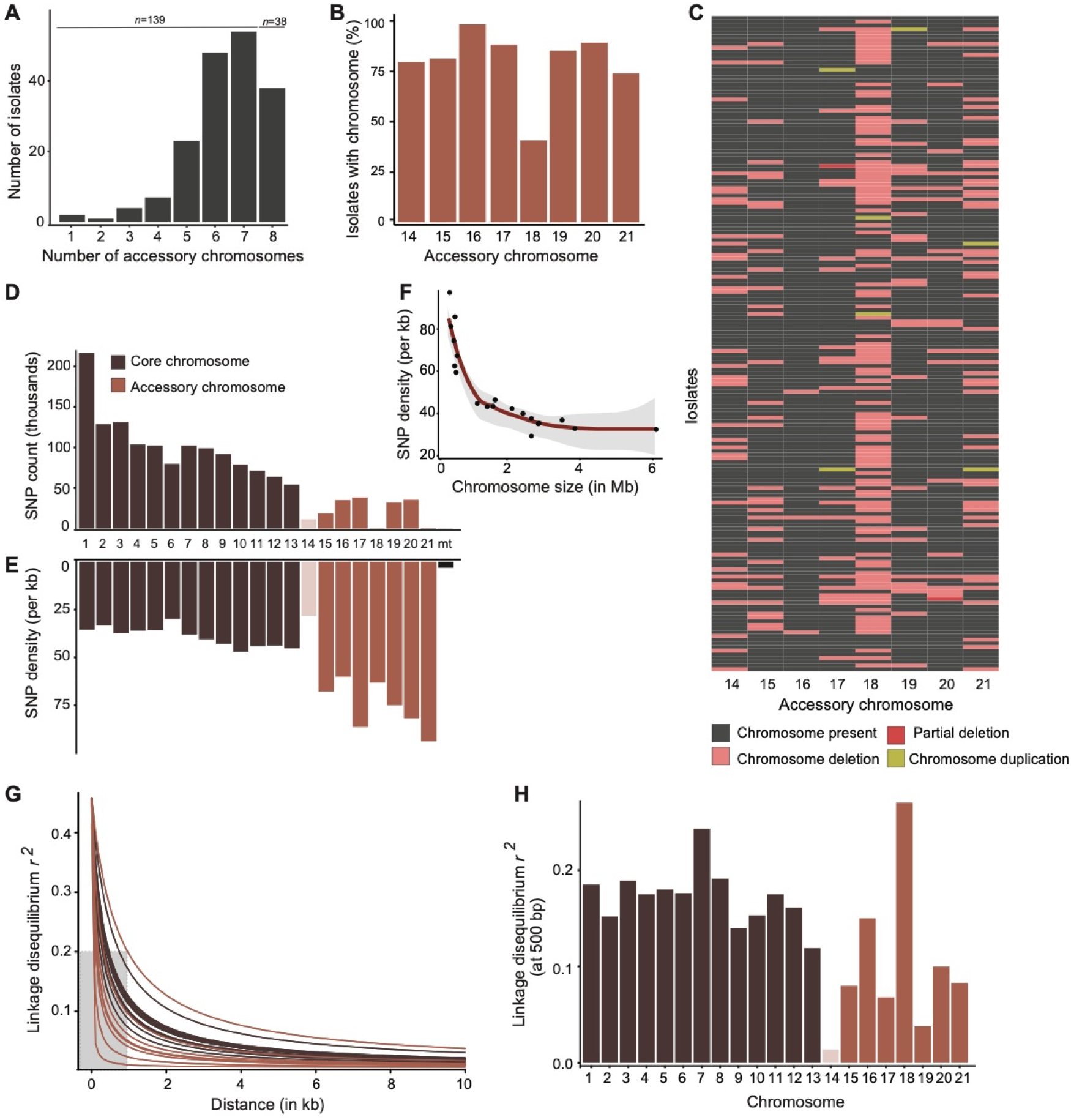
Genome-wide polymorphism and accessory chromosomes. A) Total number of accessory chromosomes per isolate. The numbers above the bars indicate the isolates missing at least one accessory chromosome or carry all chromosomes. B) Percentage presence of all the accessory chromosomes within the population calculated on the basis of normalized chromosome coverage. C) Heat map showing the presence-absence variation in accessory chromosomes across isolates assessed by normalized read depth. D) SNP count and E) SNP density per chromosome. F) Correlation plot of SNP density and chromosome size. G) Pairwise linkage disequilibrium decay among all pairs of SNPs within a fixed window size of 10’000 bp for each chromosome. A non-linear model was fitted using the equation of Ingvarsson (2005). The grey shading indicates the maximum distance needed for all chromosomes to reach *r^2^* = 0.2. H) Linkage disequilibrium *r^2^* for each chromosome at 500 bp. Light shade: chromosome 14 was omitted from per chromosome genetic diversity analyses due to the heterogeneous distribution of regions with robust SNP calls.

### Population clonality in space and time

To track the evolution of genotypic diversity in space and time, we first constructed an unrooted phylogenetic network based on SplitsTree and found that most pairs of genotypes are at a similar genetic distance (Fig 3A). The star-like structure of the network is consistent with frequent recombination. A subset of genotypes showed much shorter genetic distances suggesting recent ancestry or clonal reproduction. We also performed a PCA to determine the degree of differentiation in the population (Fig 3B). We detected no meaningful population subdivision except for seven isolates, which clustered into 3 groups (Fig 3B). Interestingly, all seven isolates were collected from cultivar Combin and two of the groups were from first collection (arrows in Fig 3B). To clarify how clonal reproduction impacts the genetic structure within the field, we identified groups of clonal genotypes based on pairwise genetic distances. We found that most isolates were at a relative pairwise distance of more than 0.15 where a value of 1 corresponds a genetic difference at every analyzed SNP position (Fig 3C). We defined isolate pairs with a genetic distance of <0.01 as being clones (Fig 3D). We assigned groups of clones into four categories according to the distance between field plots and collection time point: Clones originating from the same plot and same collection time point (category A), different plot but same collection time point (category B), same plot but different collection time point (category C) and different plot and different collection time point (category D; Fig 4A, Supplementary table S2).

**Figure 3:**
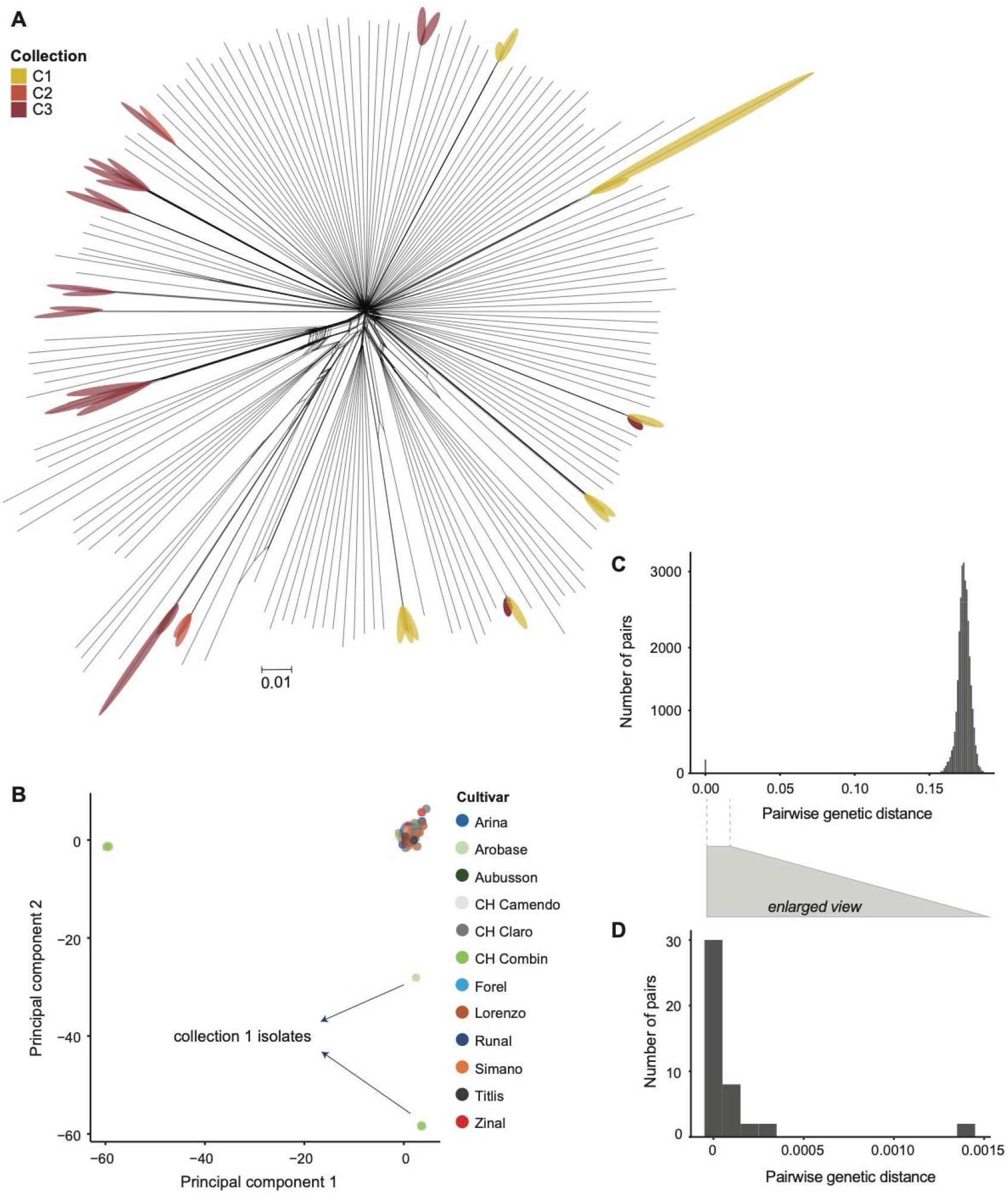
Genetic differentiation and clonality within the field. A) Phylogenetic network constructed using SplitsTree. Groups of clonal genotypes are marked with highlights identifying the collection of origin. B) The first two principal components (PC) from a PC analysis. Isolates are color coded by the cultivar of origin. C) Histograms showing the distribution of pairwise genetic distances among genotypes. A distance of 1 corresponds to the total number of SNPs. D) Histogram of pairwise genetic distances < 0.01.

**Figure 4:**
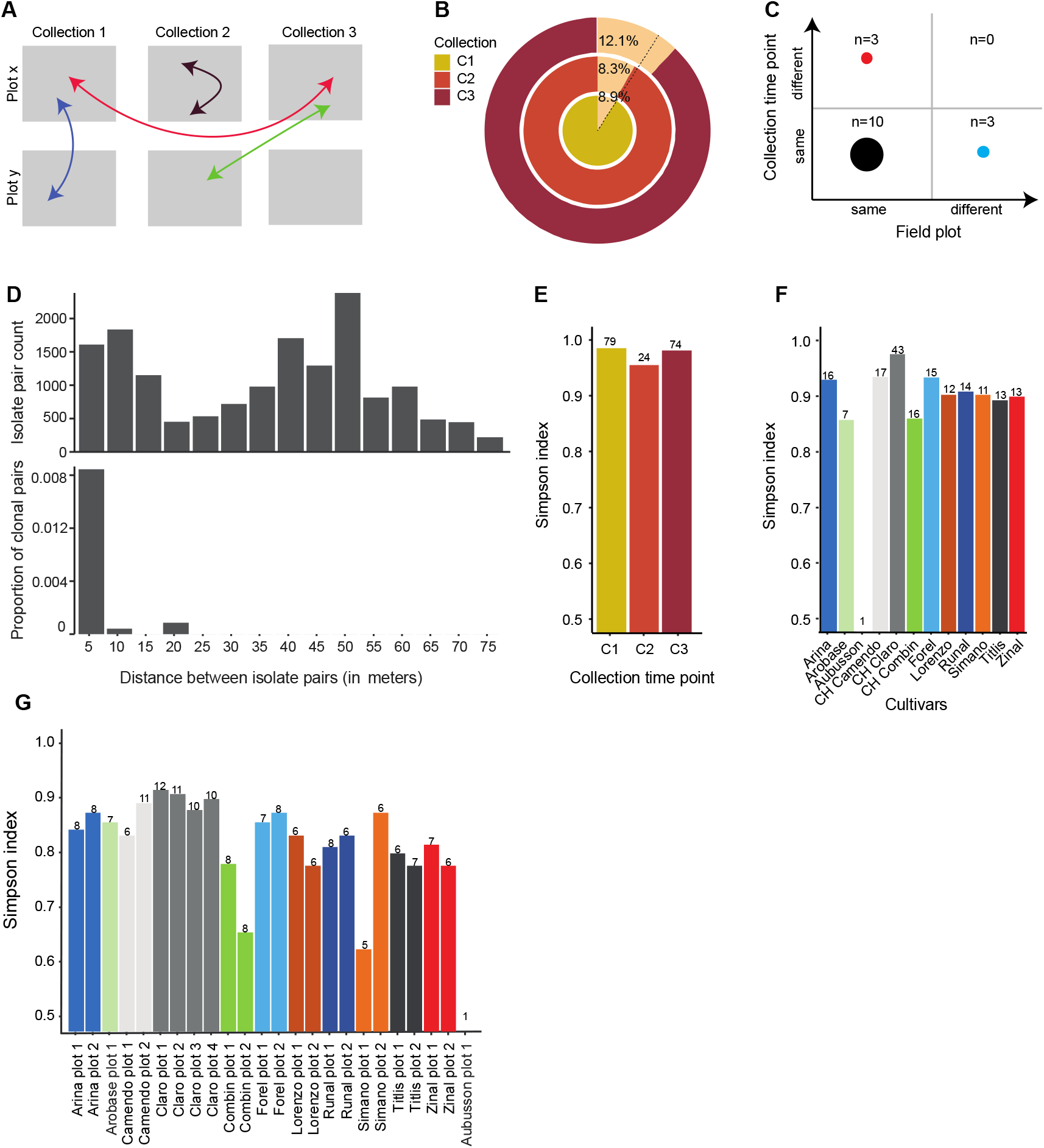
Evolution of clonality in space and time. A) Schematic representation of different clone categories with pairs identified from the same or different plots and time points. B) Pie chart showing the change in clonality over time. The dotted line indicates the average clonality of the whole population. C) Number of clone groups categorized based on differences in space and time. The numbers indicate the clone groups in each category. D) Analyses of all possible pairwise comparisons of isolates with the distance between collection points shown as a histogram. The proportion of clone pairs is shown as a proportion of the total number of isolate pairs. E) Simpson diversity index between collections C1-C3, F) between cultivars and G) plots.

We identified a total of 15 distinct groups of clones (~8.5% of the isolates; Fig 4B; Supplementary table S2). The proportion of clones ranged 8.9% in first collection to 12.1% in third collection but the difference was not statistically significant (Fisher exact test, *p-value =* 0.244; Fig 4B). The majority of the clone groups (*n* = 10) were sampled from the same plot and same collection (category A; Fig 4C-D; Supplementary table S2). Interestingly, isolates collected from the cultivar Combin showed a high degree of clonality (Supplementary table S2). We also found that 3 clonal groups each were either collected from the same plot at different time points or different plots at the same time point (category B and C; Fig 4C). The maximum distance at which we identified clonal pairs was 20 meters (Supplementary table S2) suggesting that splash dispersal can be highly effective in dispersing genotypes across the field.

We then analyzed the genotypic diversity in the field across time points using diversity indices. Overall, we found less diversity in the third collection compared to the first collection (Fig 4E Supplementary Table S3). The second collection contains too few isolates for comparison. The genotypic diversity was highest for cultivar Claro (0.97, Fig 4F) constituting also the best sampled cultivar in the field. Isolates from cultivar Combin had the lowest diversity (0.86, Fig 4F) due to the fact that almost all recovered genotypes were clones. Both field plots cultivar Combin showed low diversity with a Simpson index of 0.78 and 0.66, respectively (Fig 4G). Overall, clonal genotypes are an important constituent of the genetic structure in the field. However, the exclusive asexual reproduction during the growing season does not meaningfully affect the trajectory of genetic diversity.

## Discussion

The mode of reproduction and levels of genetic diversity play an important role in the rapid evolution of plant pathogens. We analyzed a large collection of *Z. tritici* genomes obtained from a single wheat field and found that the population was highly diverse with a rapid decay in linkage disequilibrium. We found also a minor degree of clonal structure, and clone pairs were detected both across space and across sampling time points.

### Genome-wide levels of genetic diversity in a single field

The analyzed field population shows a dominant sexual reproduction regime and is highly polymorphic. Using hierarchical sampling from the same field plots and wheat cultivars across different time points, we found that ~80% of the isolates were genetically distinct. The allele frequency spectrum and the rapid decay in linkage disequilibrium suggest that there has been no obvious recent population bottleneck. Linkage disequilibrium decayed to *r*^2^ < 0.2 within 1.5 kb across all chromosomes and often at a much shorter distance. Since the average distance between genes in the genome of *Z. tritici* is approximately 1 kb (Goodwin *et al.*, 2011), polymorphism in any gene is expected to evolve largely independently. Hence, polymorphism in genes playing roles in adaptation to either host resistance or environmental adaptation should respond independently to selection pressures allowing the population to adapt very rapidly (Slatkin, 2008; Barton, 2010). We have also shown that the rate of linkage disequilibrium decay is not the same in all chromosomes with accessory chromosomes showing faster decays. In *Z. tritici*, accessory chromosomes are small, low in G/C and gene content and have higher degrees of repetitive DNA (Goodwin *et al.*, 2011). In line with this, accessory chromosomes have been shown to have higher recombination rates than core chromosomes (Croll *et al.*, 2015). Hence, it is surprising to find the opposite trend for linkage disequilibrium in a population. A possible explanation is that the low degree of synteny among accessory chromosomes restricted genotyping to small regions, which may have biased chromosome-wide estimates of linkage disequilibrium. Rapid decay in linkage disequilibrium is an important property also in other rapidly evolving pathogens such as *Leptosphaeria maculans* causing blackleg disease in oilseed rape. Rapid allelic diversification of avirulence genes driven by sexual reproduction can lead to resistance breakdowns of the host (Daverdin *et al.*, 2012). Similarly, the poplar rust *Melampsora larici-populina* shows signs of rapid sweeps of virulent alleles associated with population replacements (Persoons *et al.*, 2017).

### Evolution of clonal genotype contributions in space and time

Sexual ascospores originating from crop residues of the previous cropping season are thought to be the source of primary infections in the field (Chen & Mcdonald, 1996; Kerdraon *et al.*, 2019). As a result of this, a wheat field infected by *Z. tritici* is expected to carry essentially no clonal genotypes at its initial colonization stage. Clonal genotypes are expected to increase in frequency as a result of splash-dispersed asexual pycnidiospores. However, we found no significant increase in clonality despite exclusive sexual reproduction. In addition to the sampling period, we found that the likelihood to observe clonal genotypes did not meaningfully change with the cultivar or physical distance. This means that even at the level of individual plots of ~2 m^2^, wheat was colonised nearly entirely by genetically distinct isolates. Notable exceptions included the clonal genotypes associated with the cultivar CH Combin. Interestingly, we identified clonal genotypes on different cultivars physically separated by as much as 20 meters. This suggests that splash dispersal has been highly effective in spreading individual genotypes across the field. Alternatively, human movement within the field may have contributed to the unusual dispersal event. We also identified clone groups with isolates collected in the same plot but from different collection time points. Hence, successful genotypes are able to persist in individual field plots at high enough levels through time to be resampled. It is currently unknown how adaptation to individual host genotypes (*i.e.* cultivars) influences the evolution of genetic diversity in infected fields. Genotypes carrying beneficial alleles to overcome host immunity should increase in frequency. In the absence of sexual recombination during the growing season, this should lead to the expansion of successful clonal genotypes. We found overall no signature of clonal expansion except for clones associated with cultivar Combin. The expansion of clones on Combin may indeed be the result of highly successful genotypes rising in frequency but population bottlenecks at the individual plot level may also produce similar patterns. Matching changes in genotype frequencies with changes in frequencies of beneficial alleles for host colonization over the course of a growing season will help to disentangle neutral from selective processes. The high degree of genotypic diversity originating from sexual reproduction is consistent with recent studies based on microsatellite markers showing spatial genetic structure but no meaningful loss in diversity over consecutive growing seasons (Siah *et al.*, 2018) (Hassine *et al.*, 2019).

Uniformity in host genotypes in agricultural ecosystems enables clonal pathogen populations to establish infections (Drenth *et al.*, 2019). However, the introduction of new host resistance genes can trigger the local extinction of such pathogen populations (Möller & Stukenbrock, 2017). Similarly, the application of fungicides can lead to local extinctions of clonal populations if populations lack resistance mutations. In contrast, in sexually reproducing populations recombination produces a multitude of new genotypes increasing the likelihood for adaptive traits to evolve (Burdon & Silk, 1997). Managing virulence and fungicide resistance emergence in pathogens has become a pressing issue to ensure sustainable agricultural production (Fisher *et al.*, 2012). Hence, understanding how pathogen populations respond in the short term (*i.e.* over the scale of weeks during the growing season) to challenges becomes a critical area of investigation. Detecting changes in pathogen populations *in situ* may become a powerful approach to identify the emergence of previously unknown adaptive mutations, changes in reproductive modes or the introduction of foreign genotypes. Deep population sequencing approaches revealing rapid changes in allele frequencies and population structure will become key tools in such endeavours.

## Materials and methods

### Field collection and storage

*Z. tritici* isolates were collected from the Field Phenotyping Platform (FIP) site of the ETH Zurich, Switzerland (coordinates 47.449°N, 198 8.682°E). A total of 335 European winter wheat varieties were grown in two replicates (Fig 1B) during the 2015–16 growing season separated each by approximately 100 m. During the collection season in 2016 the following fungicide applications were made: 4-8 April (300 g/l Spiroxamin, 160 g/l Prothioconazole), 25 May (Aviator Xpro with 75 g/l Bixafen + 150 g/l Prothioconazol), 6 June (Osiris with 56.25 g/l epoxiconazole and 41.25 g/l metconazole). Wheat was also subjected to regular applications of fertilizers and herbicides. A total of 177 isolates of *Z. tritici* were isolated from 12 winter wheat cultivars that are commonly grown in Switzerland (Levy et al., 2017) over three collection time points (fig 1A). To isolate strains, pycnidia from an infected leaf were streak-plated on a yeast sucrose broth (YSB) solid media plate and incubated at 18^°^ C for a week. Media were supplemented with 50 μg/ml kanamycin to prevent bacterial growth. After the incubation period, a single colony from the media plate was inoculated into 35 ml YSB liquid media and incubated on a shaking incubator at 18^°^C for 8 days (140-180 rpm). After the incubation period, a dense culture of blastospores was obtained. The culture was washed twice with sterile distilled water. The spores were stored as both glycerol or silica stock for future use. For the preparation of glycerol stocks, equal amounts of washed blastospores and sterile glycerol were mixed in a cryotube and stored at −80 ^°^C for future use. For the preparation of silica stocks, 200 μl of washed blastospores were poured onto 1 ml of sterile silica powder in a cryotube. Silica stocks were stored at 80 ^°^C.

### Culture preparation

Isolates were revived from glycerol stock by adding 50 μl stock solution to a 50 ml conical flask containing 35 ml liquid YSB medium. The inoculated flasks were incubated in the dark at 18^°^ C and 140-180 rpm. After 8 days of incubation, the cultures were passed through four layers of meshed cheesecloth and washed twice with sterile water to remove media traces. The filtering step also largely eliminated hyphal biomass but retained spores.

### Whole-genome sequencing and variant calling

Approximately 100 mg of lyophilized spores were used to extract high-quality genomic DNA using the Qiagen DNeasy Plant Mini Kit according to the manufacturer’s protocol. We sequenced paired-end reads of 100 bp each with an insert size of ~550 bp on the Illumina HiSeq 4000 platform. Raw reads are available on NCBI Short Read Archive under the BioProject PRJNA596434 (Oggenfuss *et al.*, 2020). We performed sequencing quality checks using FastQC v. 0.11.9. (Andrews S., 2010. www.bioinformatics.babraham.ac.uk/projects/fastqc) and extracted read counts. Sequencing reads were then trimmed for adapter sequences and sequencing quality using Trimmomatic v0.39 (Bolger et al., 2014) using the following settings: illuminaclip=TruSeq3-PE.fa:2:30:10, leading=10, trailing = 10, sliding-window = 5:10 and minlen = 50. Trimmed sequencing reads were aligned to the reference genome IPO323 (Goodwin et al., 2011) using Bowtie2 v. 2.4.1 (Langmead & Salzberg, 2012). Multi-sample joint variant calling was performed using the HaplotypeCaller and GenotypeGVCF tools of the GATK package version 4.0.1.2 (McKenna et al., 2010). We retained only SNP variants (excluding indels) and proceeded to hard filtered using the GATK VariantFiltration tool based on the following cutoffs: QD < 5.0; QUAL < 1000.0; MQ < 20.0; −2 > ReadPosRankSum > 2.0; −2 > MQRankSum > 2.0; −2 > BaseQRankSum > 2.0. After filtering for locus level genotyping rate (>80%) and minor allele count (MAC) of 1 using VCFtools v. 0.1.15 (Danecek et al., 2011). We analyzed genotyping rate at the isolate level using the --missing function of PLINK version 1.07 (Purcell et al., 2007). PLINK analyses were possible only for bi-allelic sites.

### Population genetic analyses

We performed a down-sampling analysis to estimate the effect of sample size on the total number of segregating SNPs in the population. We randomly created subsets of 177 isolates from the population and counted the number of segregating SNPs in the subset. Presence-absence polymorphism of accessory chromosomes was analyzed based on chromosome-level coverage using BAMstats v. 1.25 (https://sourceforge.net/projects/bamstats/). We calculated the standardized coverage of all chromosomes by normalizing to the mean coverage of the core chromosomes. We categorized a chromosome as present in an isolate if the standardized coverage was within the range of 0.625-1.5. Otherwise, the chromosome was categorized as partially deleted (coverage of 0.25-0.625), absent (coverage <0.25) or duplicated (coverage >1.5). We estimated SNP density on each accessory chromosome by adjusting the “--max-missing” filter for SNPs to 80% of the total number of observed chromosomes. We used the loess model implemented in the geom_smooth function from the R package ggplot2 (Wickham, 2016).

### Linkage disequilibrium and population structure analyses

We analyzed the decay in linkage disequilibrium for each chromosome separately. For this, we used all SNPs with a minor allele frequency >5% in the population. We calculated the linkage disequilibrium *r*^2^ between marker pairs using the option–hap-r2 in VCFtools v. 0.1.15 (Danecek *et al.*, 2011) with --ld-window-bp of 10000. The decay of linkage disequilibrium with physical distance was estimated using a nonlinear regression model (Remington *et al.*, 2001; Ingvarsson, 2005). We performed and visualised principal component analyses (PCA) using the R packages vcfR v. 1.8.0 (Knaus & Grünwald, 2017), adegenet v. 2.1.1 (Jombart & Ahmed, 2011), and ggplot2 v. 3.1.0 (Wickham, 2016). We generated an unrooted phylogenetic network using SplitsTree v4.14.6 (Huson, 1998). File format conversions were performed using PGDSpider v2.1.1.5 (Lischer & Excoler, 2011). To identify groups of clonal genotypes, we calculated the pairwise genetic distances between all genotypes using the function dist.dna() included in the R package ape v. 5.3 (Paradis & Schliep, 2019). Isolate pairs with a pairwise genetic distance below 0.01 were considered as clones for further analyses. The Simpson diversity index was calculated using the R package vegan v. 2.5 (Oksanen *et al.*, 2019).

## Acknowledgement

Luzia Stalder and Ursula Oggenfuss provided helpful comments on a previous version of the manuscript. Petteri Karisto performed the field sampling and established the strain collection. Leen Abraham provided assistance in the laboratory. The DNA library for the sequencing was prepared at the Genetic Diversity Centre (GDC) facility of ETH, Zurich. Sequencing was performed at the iGE3 platform of the University of Geneva. EC was supported by the European Union’s Horizon 2020 programme under the Marie Skłodowska-Curie grant agreement No 799630 (PATH2EVOL). DC is supported by the Swiss National Science Foundation (grants 31003A_173265 and IZCOZO_177052).

