## Supplementary Figure 1 for "Population-level deep sequencing reveals the interplay of clonal and sexual reproduction in the fungal wheat pathogen *Zymoseptoria tritici*"

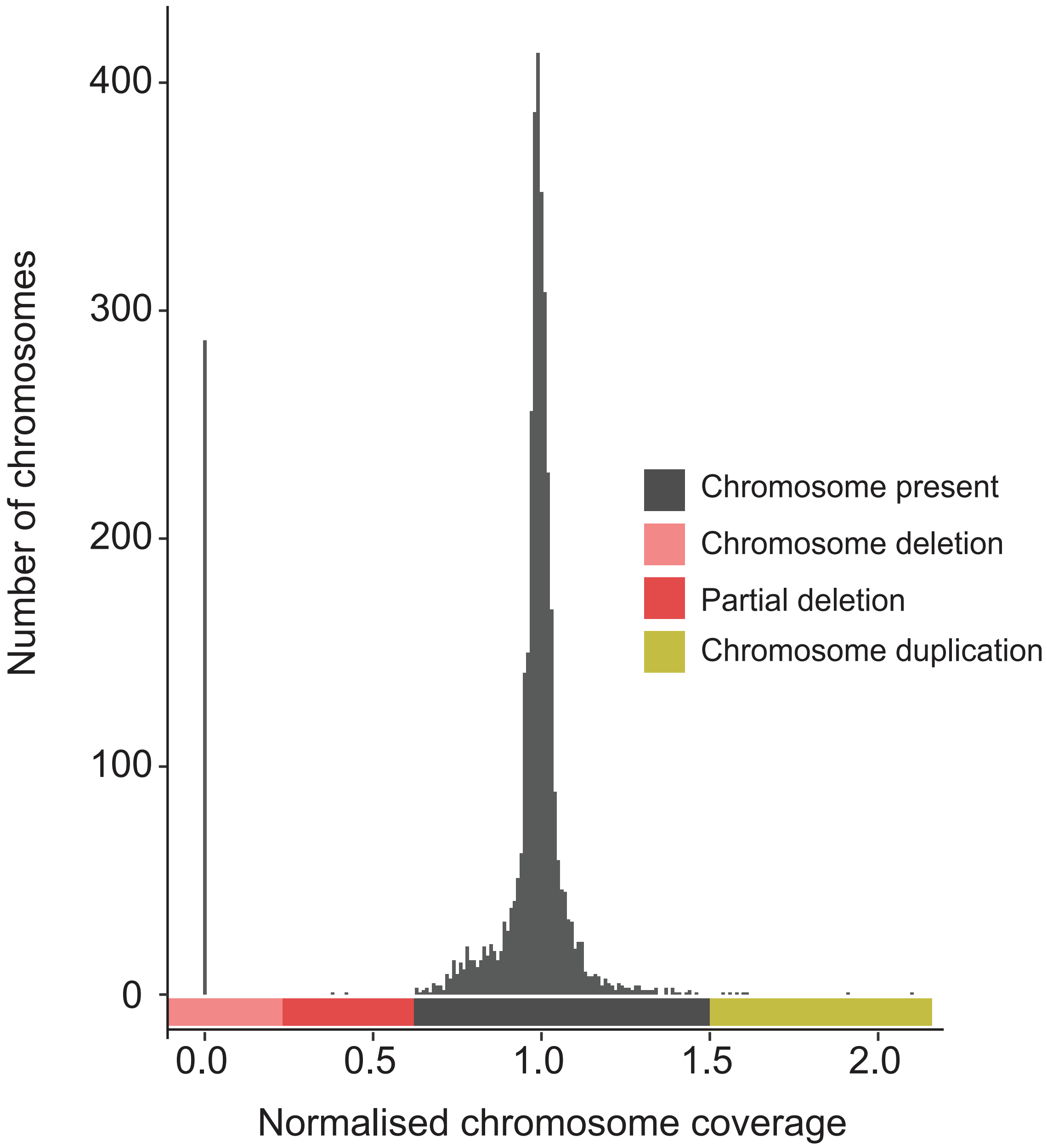


**Supplementary Figure S1**: Normalized chromosome coverage of core and accessory chromosomes. The colored bars indicate normalized coverage cutoffs for defining the presence, absence, partial deletion and duplication state of individual chromosomes.
